# High bacillary burden and the ESX-1 type VII secretion system promote MHC class I presentation by *Mycobacterium tuberculosis* infected macrophages to CD8 T-cells

**DOI:** 10.1101/2022.10.20.513087

**Authors:** Daniel Mott, Jason Yang, Christina Baer, Kadamba Papavinasasundaram, Christopher M. Sassetti, Samuel M. Behar

**Author notes:** Correspondence: Samuel M. Behar, E-mail address (SMB).

## Abstract

*Mycobacterium tuberculosis* (Mtb) subverts host defenses to persist in macrophages despite immune pressure. CD4 T-cells can recognize macrophages infected with a single bacillus *in vitro*. Under identical conditions, CD8 T-cells inefficiently recognize infected macrophages and fail to restrict Mtb growth, although they can inhibit Mtb growth during high burden intracellular infection. We show that high intracellular Mtb numbers cause macrophage death, leading other macrophages to scavenge cellular debris and cross-present the TB10.4 antigen to CD8 T-cells. Presentation by infected macrophages requires Mtb to have a functional ESX-1 type VII secretion system. These data indicate that phagosomal membrane damage and cell death promote class I MHC presentation of the immunodominant antigen TB10.4 by macrophages. Although this mode of antigen-presentation stimulates cytokine production that we presume would be host beneficial; killing of uninfected cells could worsen immunopathology. We suggest that shifting the focus of CD8 T-cell recognition to uninfected macrophages would limit the interaction of CD8 T-cells with infected macrophages and impair CD8 T-cell mediated resolution of tuberculosis.

## Introduction

Despite concerted efforts to eradicate tuberculosis (TB), it persists as one the deadliest global infectious diseases^1^. First tested in 1921, Bacillus Calmette–Guérin (BCG) remains the only approved vaccine against *Mycobacterium tuberculosis* (Mtb), and yet its efficacy is variable among human populations, and it fares poorly at preventing pulmonary tuberculosis^2,3^. Impaired cellular immunity caused by conditions as diverse as malnutrition and HIV infection are risk factors for progression to active TB, which highlight the importance of T-cells in containing Mtb infection^4–6^. As in humans, T-cell responses are essential for control of Mtb infection in mice^7,8^. Although CD4 T-cells appear to contribute more to protection than CD8 T-cells in mice, optimal protection is achieved only when both T-cell subsets are present^9,10^. The success of Mtb as a pathogen is attributed to its immune evasion strategies, including manipulation of vesicular trafficking and exploitation of immune responses to generate favorable environments for bacterial growth, dissemination, and transmission^11,12^. With an increased incidence of TB caused by multi-drug resistant strains, the push for a more effective vaccine is urgent. To develop more effective vaccines, we need a better understanding of protective immunity^1^.

Viral and intracellular microbes such as influenza and *Listeria monocytogenes* assemble or replicate in the cytosol of infected cells. There, their antigens are sampled by class I MHC and presented to CD8 T-cells, which play a key role in protection against these pathogens^13,14^. Canonical MHC I presentation samples cytosolic antigens, which are processed by the proteasome and loaded onto MHC I in the endoplasmic reticulum (ER)^15^. For extracellular antigen sources (e.g., extracellular bacteria or tumor cells), antigen presenting cells (APC) engulf the pathogen or apoptotic tumor cell by phagocytosis or efferocytosis, respectively, and use cross-presentation to shuttle antigens from endosomal compartments into the MHC I presentation pathway^16^. These pathways are crucial for the presentation of intracellular pathogens that reside in endosomal compartments. While cross-presentation is associated with DC priming of T-cells, macrophages also cross-present antigens to CD8 T-cells ^17^. As most Mtb-infected cells are macrophages^18^, we are interested in how macrophages present Mtb antigens to CD8 T-cells.

Mice mount a substantial CD8 T-cell response following Mtb infection. In infected C57BL/6 (B6) mice, up to 40% of CD8 T-cells in the lung recognize the TB10.4_4-11_ epitope, making TB10.4 the most immunodominant antigen in this model^19^. To determine the potential of vaccine-elicited CD8 T-cells to mediate protection against Mtb infection, we generated TB10.4_4-11_-specific memory CD8 T-cells using a peptide-based vaccine; however, TB10.4_4-11_-specific CD8 T-cells failed to enhance control of Mtb challenge^20^. In parallel, we generated TB10.4_4-11_-specific CD8 T-cell lines, which express TCRs of different affinities (e.g., Rg3 and Rg4)^21^. The Rg3 and Rg4 CD8 T-cell lines are activated when they recognize their cognate antigen either in the form of γ-irradiated Mtb or synthetic peptide, but they fail to recognize lung myeloid cells isolated from Mtb-infected mice, or *in vitro* infected BMDM or thioglycolate-elicited peritoneal macrophages (TGPM)^22,23^. Under identical conditions and in stark contrast, CD4 T-cell lines specific for the Mtb antigens Ag85B or ESAT6 recognize Mtb-infected macrophages and inhibit intracellular Mtb growth^20,23^.

We suggest that the poor ability of vaccine-elicited TB10.4-specific CD8 T-cells to protect mice against Mtb arises from their inability to recognize Mtb-infected macrophages despite their efficient priming and persistence during infection. The poor recognition of infected macrophages could explain why CD8 T-cells have only a modest impact on control of Mtb infection. The discrepancy between the immunodominance of TB10.4-specific CD8 T-cells and their inability to recognize Mtb-infected macrophages, is paradoxical and has led us to hypothesize that TB10.4 is a decoy antigen^22,23^. These data motivated us to determine under what conditions Mtb antigens are presented by macrophages and recognized by CD8 T-cells.

## Results

### Infected macrophages poorly activate Mtb-specific CD8 T-cells

To determine whether the inability of TB10.4_4-11_-specific CD8 T-cells to recognize infected macrophages is idiosyncratic to the TB10.4_4-11_ epitope and B6 mice, we determined whether CD8 T-cells specific for other Mtb epitopes could recognize Mtb-infected macrophages. The Mtb32A protein (i.e., Rv0125) is a second well-described antigen recognized by CD8 T-cells ^24,25^. The CD8 T-cell response to Mtb32A_309-318_ is subdominant to TB10.4_4-11_-specific CD8 T-cells, but accounts for a significant fraction of lung CD8 T-cells in the lung early after infection. A polyclonal CD8 T-cell line was generated from Mtb32A_309-318_ peptide-vaccinated B6 mice and 90% of the CD8 T-cell line stained with the K^b^/Mtb32A_309-318_ tetramer. The Mtb32A-specific CD8 T-cell line produced IFN-γ when cultured with macrophages pulsed with Mtb32A_309-318_ peptide or with γ-irradiated Mtb [Fig.1A, left]. However, like TB10.4_4-11_-specific CD8 T-cells, it failed to recognize to Mtb-infected macrophages. A second CD8 T-cell line (i.e., DM108) was generated and is specific for TB10.4_20-28_, which is an immunodominant epitope in Mtb-infected BALB/c mice and presented by H-2 K^d^. The DM108 CD8 T-cell line produced IFN-γ when cultured with macrophages pulsed with TB10.4_20-28_ peptide or γ-irradiated Mtb, but not Mtb-infected macrophages [Fig.1B, right]. Including the TB10.4_4-11_ epitope, these data show that three different epitopes from two different proteins in two different mouse genetic backgrounds are inefficiently presented by Mtb-infected macrophages to class I MHC-restricted CD8 T cells.

**Figure 1.**
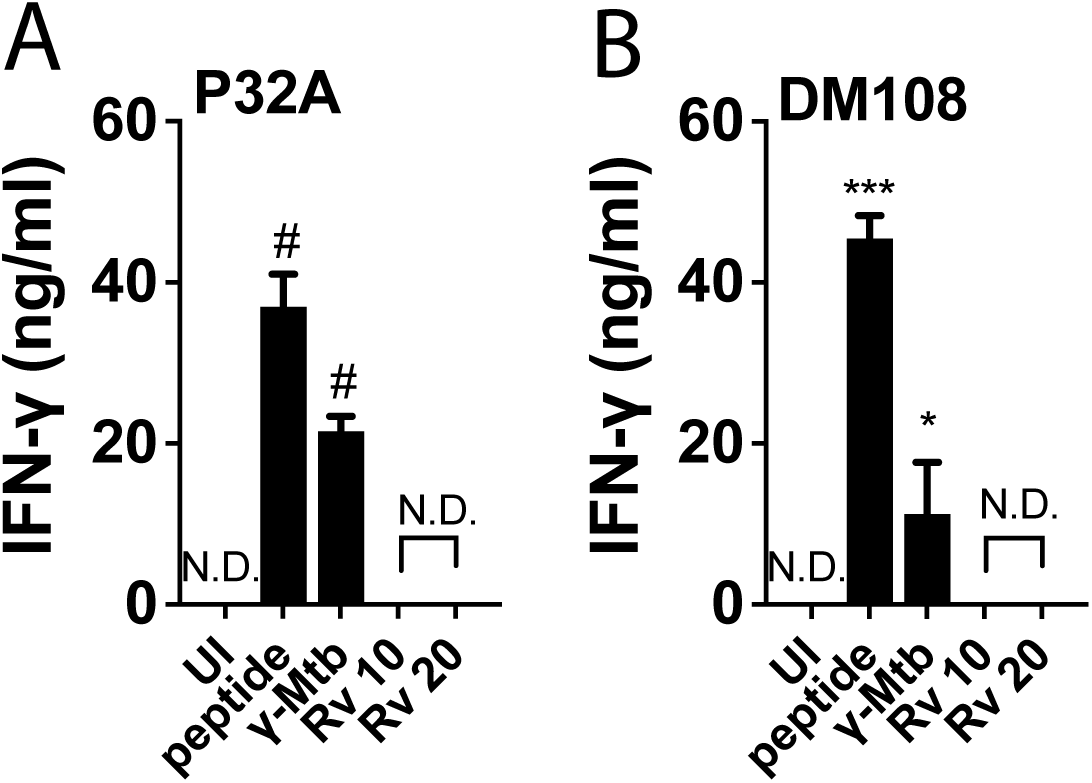
Infected macrophages poorly activate Mtb-specific CD8 T-cells. Polyclonal CD8 T-cell lines specific for Mtb32A_309-318_ (i.e., P32A) (A) or TB10.4_20-28_ (i.e., DM108) (B). CD8 T-cell line recognition of TGPM that were uninfected (UI), pulsed with cognate peptide (peptide), γ-irradiated Mtb (γ-Mtb) or live Mtb H37Rv infection at initial MOIs of 10 or 20 (Rv 10, Rv 20). Bar, mean ± SD. The data are representative of at least two independent experiments. Statistical test: one-way ANOVA, using the Dunnett test compared to uninfected macrophages. P values: *, p<0.05; #, p<0.0001. N.D., none detected.

### Infected macrophages cross-present antigen but fail to activate Mtb-specific CD8 T-cells

The poor recognition of infected macrophages by Mtb-specific CD8 T-cells raises the possibility that infected macrophages fail to cross-present phagosomal antigens via class I MHC. While cross-presentation is best characterized in DC, macrophages cross-present antigen as well^17^. To assess the ability of TGPM to cross-present antigen, TB10.4 protein-coated beads were added to TGPM and then cultured with TB10.4-specific CD8 T-cells. TB10.4-bead-treated TGPM activated the Rg3 TB10.4-specific CD8 T-cell line^21,23^, in a dose dependent manner, verifying the capacity of TGPM to cross-present antigen [Fig.2A].

**Figure 2.**
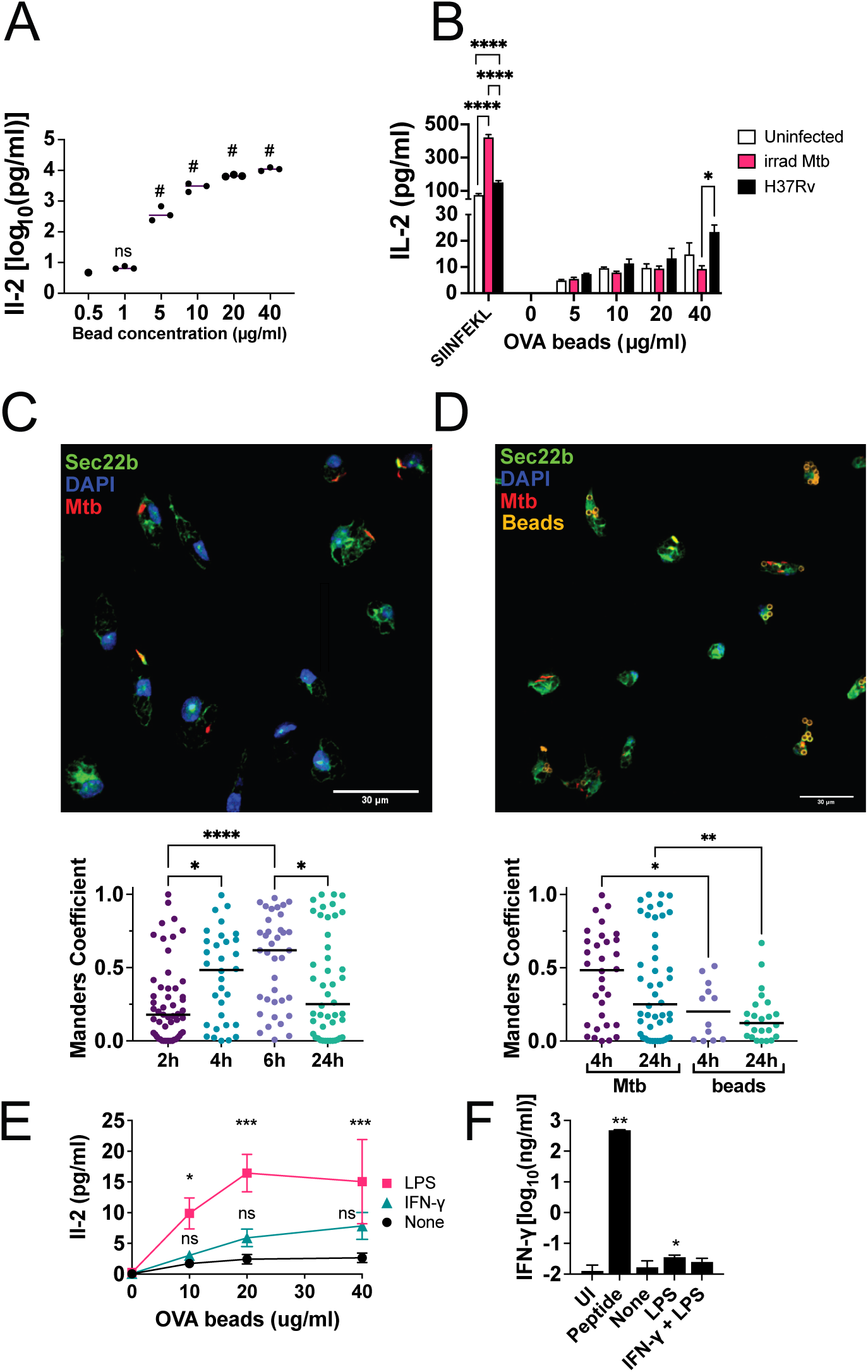
Infected macrophages cross-present antigen but fail to activate Mtb-specific CD8 T-cells. (A) Rg3 CD8 T-cells recognized TB10.4 protein-coated beads presented by TGPM. (B) RF33.70, an OVA-specific MHC I restricted CD8 hybridoma, recognized OVA-coated beads presented by uninfected, γ-irradiated Mtb, or H37Rv-infected TGPM. (C, D) Top. Representative images of H37Rv.YFP infected-TGPM stained for Sec22b without (C) or with (D) latex beads. Bottom. Mander’s coefficient of Sec22b colocalization with Mtb and latex beads at the indicated time-points. Each point represents an individual Mtb-infected macrophage. (E) Effects of LPS or IFN-γ treatment on cross-presentation of OVA-bead to OT-I CD8 T-cells. (F) Rg4 CD8 T-cell recognition of H37Rv-infected B6 TGPM pre-treated with LPS or LPS+IFN-γ. Bars, mean ± SD. The data are representative of three (B) or two (C - F) independent experiments, (n=2-3 replicates/condition). Statistical analysis: one-way ANOVA of log_10_ transformed data using Dunnett’s multiple comparison test compared to 0.5 ug/ml TB10.4-beads (A), two-way ANOVA with Fisher’s LSD posttest (B), or by one-way ANOVA with Fisher’s LSD test (C - F) p-values: *, p<0.05; **, p<0.01; ***, p<0.001; #, p<0.0001.

TGPM cross-present TB10.4 to CD8 T-cells, whether the antigen is coated onto beads [Fig.2A] or in the form of γ-irradiated Mtb^23^. TGPM infected with TB10.4-expressing *Listeria monocytogenes* also present the TB10.4_4-11_ epitope to Rg3 CD8 T-cells, by a listerolysin dependent pathway. As macrophages can present TB10.4 when it is provided in different forms, we hypothesized that live Mtb infection globally inhibits cross-presentation. If true, we reasoned that cross-presentation of ovalbumin (OVA) would be impeded in Mtb-infected macrophages. Uninfected macrophages or macrophages infected with Mtb or fed γ-irradiated Mtb were pulsed with OVA-coated beads and then cultured with the OVA_257-264_-specific RF33.70 T-cell hybridoma^26^. Mtb-infected macrophages cross-presented OVA_257-264_ to RF33.70, showing that Mtb did not interfere with the ability of TGPM to cross-present exogenous antigen [Fig.2B].

An alternative hypothesis is that Mtb infection only inhibits cross-presentation within Mtb-containing compartments, which could explain why OVA-bead presentation was unaffected. To test this possibility, we measured the recruitment of the cellular cross-presentation machinery to H37Rv.YFP-containing phagosomes. Sec22b contributes to the cytosolic pathway of cross-presentation by facilitating trafficking of ER components into phagosomes^27–29^. We found strong colocalization between Sec22b and Mtb in TGPM, which increased over time with mean Mander’s coefficient from 0.26 – 0.54 [Fig.2C]. More Sec22b colocalized with Mtb than with latex beads [Fig.2D]. We found significant colocalization of the cross-presentation markers P97^30,31^ and MHC-I, but not the vacuolar component Cathepsin S or early endosome marker Rab5 [Fig.S1]. The role of these proteins in cross-presentation has been primarily characterized in DC; nevertheless, these data show that Mtb does not interfere with their recruitment to phagosomes in macrophages.

We next tested whether macrophage activation enhances cross-presentation of Mtb antigens to T-cells. Pre-activating macrophages enhances their phagocytic and antigen presentation capacity ^32,33^, and while Mtb infection stimulates TLR signaling^34,35^, we hypothesized that activation of other pathways might enhance cross-presentation. Treating TGPM with LPS, but not IFN-γ, enhanced their presentation of OVA-coated beads to the RF33.70 CD8 T-cell hybridoma [Fig.2E]. However, neither of these treatments enhanced cross-presentation of TB10.4 by infected macrophages to Rg3 cells [Fig.2F].

DC more efficiently cross-present antigen to CD8 T-cells than macrophages^17,36^. Therefore, we tested whether Mtb infection of bone-marrow (BM) derived DC (BMDC) or HoxB8 differentiated DC ^37,38^ could activate CD8 T-cells. These DC, like the experiments with macrophages, did not activate TB10.4-specific CD8 T-cells [Fig.S2]. As the cDC1 subset most efficiently cross-presents antigen to CD8 T-cells, we used Batf3^−/−^ mice, which have a developmental deficiency of cDC1 to assess their role in priming TB10.4-specific CD8 T cells ^39^. The number of pulmonary TB10.4- and 32A-specific CD8 T-cells was similar in Mtb-infected Batf3^−/−^ and B6 mice, indicating that Batf3-dependent cDC1 are not required for priming CD8 T-cell responses to these immunodominant Mtb antigens [Fig.S3]. We suggest that a different type of APC is priming these responses, or there is compensatory development of CD8α^+^ DC during Mtb infection^40^. These results suggest that while Mtb does not globally interfere with cross-presentation in macrophages, Mtb-containing phagosomes avoid cross-presentation to CD8 T-cells under conditions that permit presentation to CD4 T-cells.

### High intracellular bacterial burden leads to cross-presentation

We recently identified conditions that an overnight infection maximized MOI and minimized cell necrosis to optimize recognition of infected macrophages by polyclonal T-cells ^10,22,23,41^ Under these optimal conditions, only 10% of polyclonal CD8 T-cells from the lungs of infected mice recognize Mtb-infected macrophages *in vitro,* based on IFN-γ production. Also, polyclonal CD4 T cells recognize Mtb-infected macrophages more efficiently (i.e., at a lower MOI) than polyclonal CD8 T cells. We revisited this issue to determine how varying the MOI overnight affects macrophage cell death and T cell recognition.

As expected, increasing the number of Mtb added to macrophages (i.e., the MOI) for 24 hours correlated with cell death [Fig.3A, left]. By determining the number of intracellular Mtb, we calculated a corrected MOI adjusted for cell death [Fig.3A, middle and right]. These results suggest that despite massive cell death, there exist populations of highly infected macrophages at the highest MOIs. Under conditions accompanied by massive cell death, we expect considerable heterogeneity in the number of Mtb per macrophage at the single cell level. We used an experimental approach to enable image analysis of all macrophages in each condition. Macrophages were infected with H37Rv.YFP for two hours or 24 hours at several different MOIs. We quantified the pixels of in each macrophage and inferred the apparent bacteria/macrophage ratio [Fig.3B]. After the standard two-hour (low MOI) infection, 16-27% of the macrophages were infected [Fig.3C]. The infected macrophages had a median of 2.0-2.3 Mtb/macrophage [Fig.3D]. Overnight (high MOI) infection led to infection of 79-86% of macrophages, with a median of 6.3-12.4 Mtb/macrophage. As expected, we found that after overnight infection with high MOIs, there was significant heterogeneity in the macrophage bacterial burden [Fig.3D].

**Figure 3.**
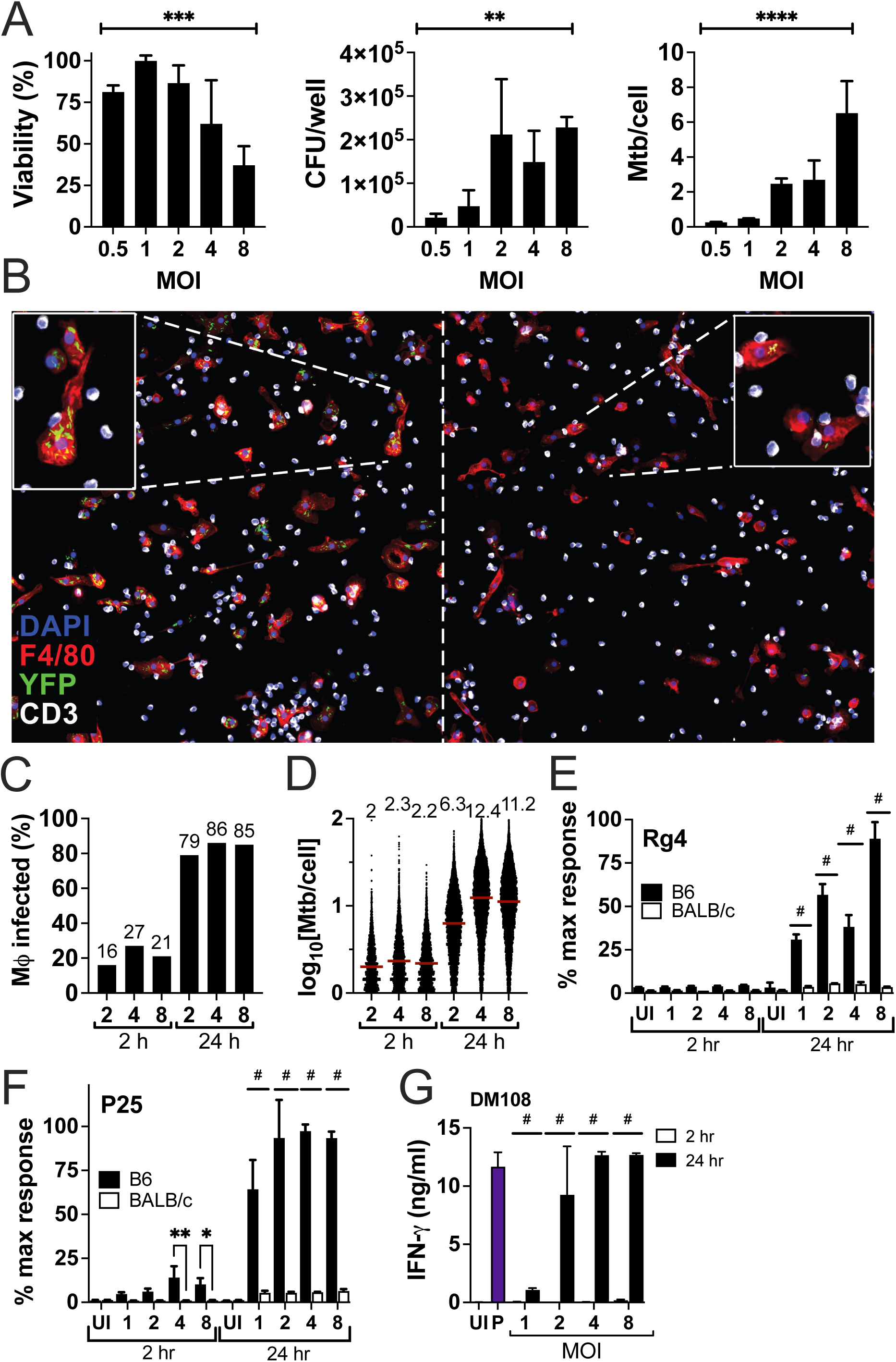
High MOI infection promotes antigen presentation by macrophages to CD8 T-cells. (A) Viability (left), intracellular CFU (middle), and adjusted MOI (intracellular CFU/number of viable cells) (right) of TGPM infected for 24-hours. (B) Representative images of 24-hour (left) and 2-hour (right) H37Rv.YFP-infected HOXB8 macrophages cultured with Rg4 CD8 T-cells. Inset, enlargement of indicated macrophages. (C, D) Image analysis of HOXB8 macrophages infected with H37Rv.YFP with different MOIs for 2- or 24-hours. For each condition, 13,000-18,000 macrophages were analyzed. (C) Percent of macrophages infected with H37Rv.YFP. (D) Distribution of bacteria/macrophage for each condition evaluated. Uninfected macrophages were excluded from analysis and the bacterial number was extrapolated as described in the *Methods*. (E) Rg4 and (F) P25 recognition of B6 [H2-matched] or BALB/c [mismatched] H37Rv-infected TGPM for 2-hours (low MOI) or 24-hours (high MOI). (G) DM108 CD8 T-cell recognition of HOXB8 macrophages infected for 2 or 24 hours with H37Rv. Bar and numbers above bar represent median. Data representative of three (A) or two (B - G) independent experiments. Bars represent mean ± SD (n=3 replicates/condition). Statistical testing: (A) Ordinary one-way ANOVA testing for a linear trend between column means; (E, F) two-way ANOVA with Šidák correction for multiple comparisons; or two-way ANOVA with Fisher’s LSD posttest (G). p values: *, p<0.05; **, p<0.01; ***, p<0.001; #, p<0.0001.

We next asked how these two different infection conditions affect antigen presentation to CD4 and CD8 T-cells. Rg4 CD8 T-cells specifically recognized B6 macrophages infected for 24 hours but not 2 hours [Fig.3E]. As reported previously, P25 CD4 T-cells did recognize low MOI infected macrophages (i.e., infected for 2 hours). The response of P25 cells was more robust when cultured with the high MOI infected macrophages [Fig.3F]. Neither Rg4 nor P25 produced IFN-γ when cultured with Mtb-infected BALB/c (i.e., H-2 mismatched) macrophages. Thus, IFN-γ production by these T cells is H-2 restricted and was not the result of noncognate activation (e.g., cytokine-mediated stimulation). Thus, both at a low and high effective MOI, Mtb-infected macrophages more efficiently presented antigen to CD4 T-cells than CD8 T-cells. We occasionally observed that IFNγ-production by CD4 T-cells, and to a lesser extent CD8 T-cells, was decreased at the highest MOI in some experiments, which we attributed to the number of viable APC falling below a critical threshold secondary to cell death (Fig.S4). Similarly, the H-2^d^-restricted TB10.4_20-28_-specific CD8 T-cell line (i.e., DM108) recognized macrophages infected at a high MOI but did not recognize macrophages infected at low MOIs [Fig.3G]. Based on these data, we hypothesized that TB10.4-specific CD8 T-cells preferentially recognize highly infected macrophages.

### Interaction of TB10.4-specific CD8 T-cells with Mtb-infected macrophages

Measuring IFNγ as the only indication of T cell recognition of infected macrophages could bias our estimates of T cell activation. Not all activated T cells produce IFNγ and not all IFNγ is the result of cognate recognition. To measure CD8 recognition of Mtb-infected macrophages independently of function, we enumerated stable interactions between CD8 T-cells and Mtb-infected macrophages. To distinguish cognate from noncognate interactions, CD8 T cell interaction with WT vs. MHC I^−/−^ macrophages was compared. We leveraged the heterogeneity of the *in vitro* infection and our capacity to enumerate thousands of cells to determine the propensity of CD8 T-cells to interact with macrophages over a range of MOIs. Few CD8 T-cells interacted with uninfected or low MOI (i.e., 1 or 2 Mtb/macrophage) infected WT or MHC I^−/−^ macrophages [Fig.4A]. However, as the MOI increased, significantly more T cells interacted with Mtb-infected WT macrophages compared to MHC I^−/−^ macrophages. This preference was observed between a range of MOI = 3-18 [Fig.4B]. The mean T-cell contacts with infected WT HOXB8 macrophages were significantly correlated with intracellular bacilli (Spearman ρ = 0.19, p<0.0001). In contrast, there was no correlation between T-cells interacting with MHC I^−/−^ macrophages and the number of intracellular Mtb (Spearman ρ = 0.0236, p=n.s.). These data confirm that T-cell-macrophage interactions are TCR-MHC dependent, and taken together, support the hypothesis that CD8 T-cells preferentially recognize highly infected macrophages and poorly recognize macrophages containing few bacilli.

**Figure 4.**
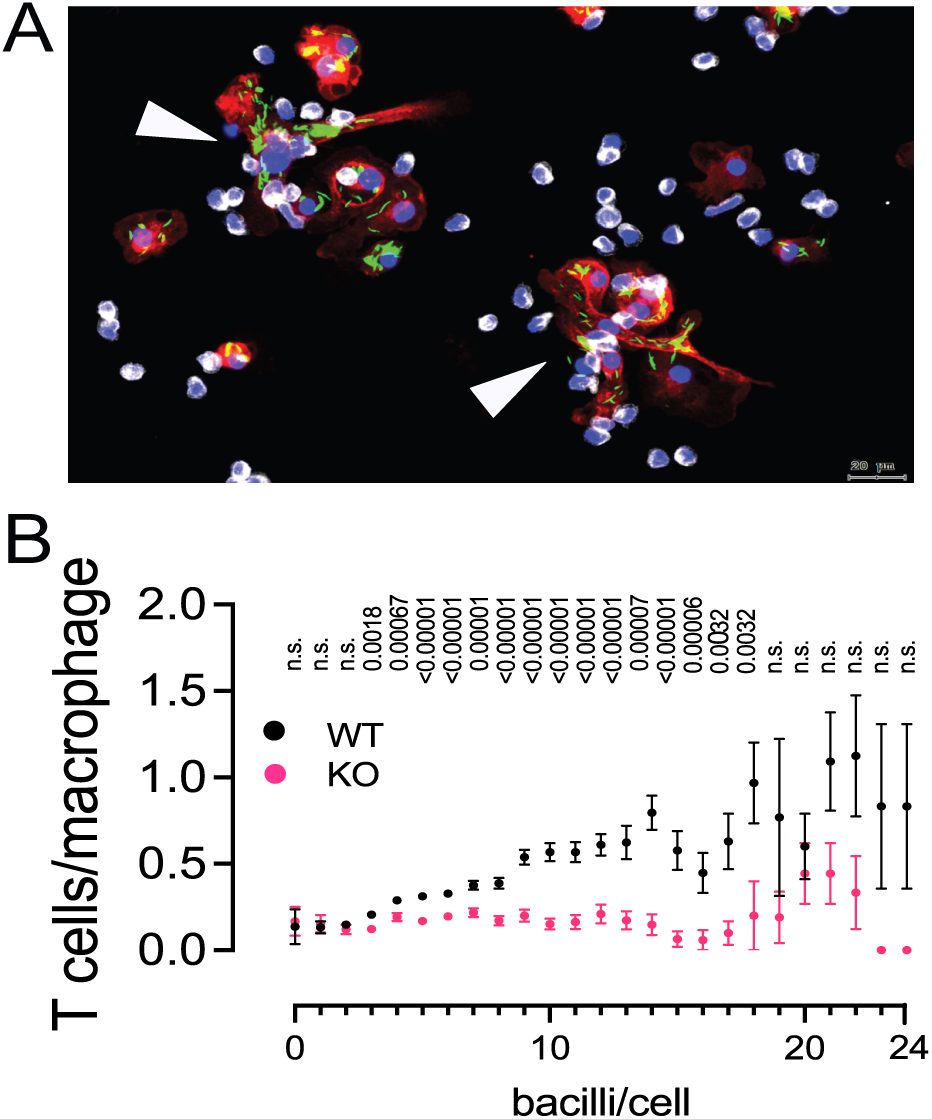
Interaction of TB10.4-specific CD8 T-cells with Mtb-infected macrophages. (A) Representative image of highly infected macrophages with many interacting T-cells (arrows, 9 and 10 interacting T-cells), and sparsely infected macrophages with no T-cell contacts. (B) Mean T-cell contacts with infected WT or MHC I^−/−^ (KO) HOXB8 macrophages as a function of intracellular bacterial content. Data representative of two independent experiments. Each point is mean ± SEM of T cells/macrophage at each inferred MOI. Multiple unpaired t test with Welch correction were performed, with a Two-stage step-up (Benjamini, Krieger, and Yekutieli) and a false discovery rate set to Q=1%. The values shown in the figure represent q values.

### Cell death correlates with antigen presentation to CD8 T cells

Why a high bacterial burden should drive MHC I presentation was not intuitively obvious as the inability of macrophages to present TB10.4 did not appear to be an issue of antigen abundance^23^. We hypothesized that a mechanistic link exists between the death of Mtb-infected macrophages and MHC I presentation of Mtb antigens ^42–47^. In vitro, Mtb-infected macrophages die primarily by necrosis^43,44,46^, which is enhanced by high bacterial burdens^47–49^, and these necrotic macrophages can be engulfed by other phagocytes^50,51^. We hypothesize that Mtb antigens from dying infected cells are acquired by phagocytic cells that secondarily present them to CD8 T-cells. To test this possibility, we used Celltracker™ dyes to differentiate between primary infected (Deep Red) and uninfected bystander (VioGreen) macrophages [Fig.5A]. After culturing infected and uninfected macrophage together, between 2-6% of the macrophages that were originally uninfected now contained H37Rv.YFP, indicating that they were secondarily infected [Fig.5B, left]. Of these secondarily infected macrophages, 75-83% were also positive for the Deep Red dye that was used to label infected macrophages [Fig.5B, right]. These data indicate that the majority of “secondarily infected” macrophages had engulfed dying infected macrophages. These findings were confirmed by immunofluorescence microscopy [Fig.5C], and among the secondarily infected macrophages (i.e., VioGreen^+^), we identified a strong correlation between H37Rv.YFP and acquisition of Deep Red dye (i.e., from the originally infected macrophages) (R^2^ = 0.4375) [Fig.5D]. We conclude that cell death driven by high MOI Mtb infections can lead to bystander macrophage phagocytosis of dead cellular debris containing Mtb.

**Figure 5.**
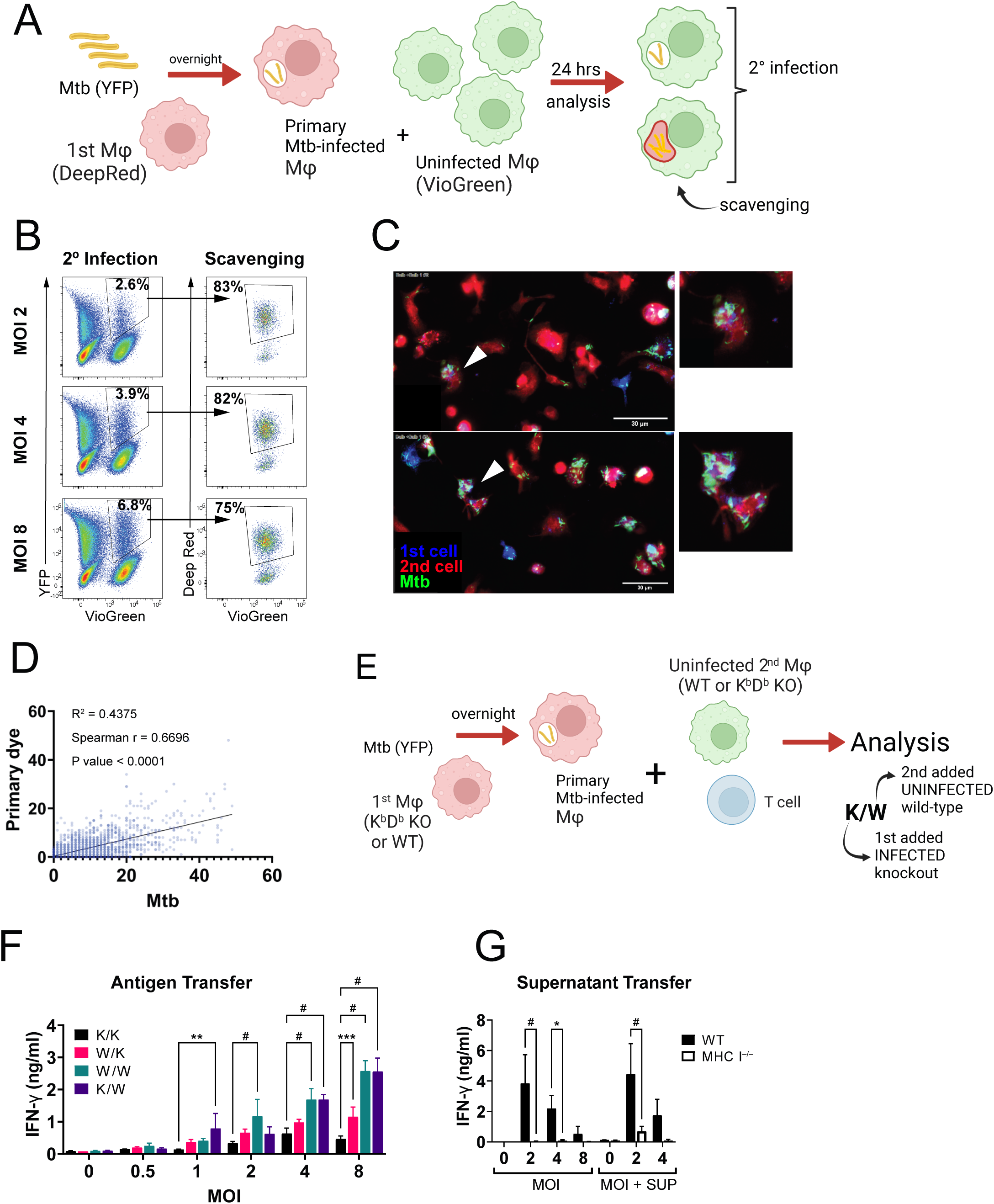
Cell death contributes to cross-presentation. (A) Schematic of scavenging assay. Primary macrophages (Mj) were infected with H37Rv.YFP overnight. Cells were washed, and secondary uninfected macrophages were added and cocultured for 24 hrs. (B) Representative flow plots following co-culture of differentially stained macrophages. Initial gating reveals the percentage of bystander macrophages (Viogreen^+^) that took up Mtb (YFP^+^) (left). The Viogreen^+^YFP^+^ macrophages were gated to determine the percentage that acquired Deep Red dye from primary-infected macrophages (right). (C) Representative image of scavenging assay. Arrows indicate bystander macrophages that acquired Mtb and dye from primary-infected macrophages. Inset, enlargement. (D) Analysis of the bystander macrophages showing that uptake of Mtb and primary-infected macrophages are correlated. (E) Schematic of antigen transfer assay. APC are represented as wild-type (“W”) or MHC I^−/−^ (“K”) HOXB8 macrophages (e.g., K/W = infected MHC I^−/−^, uninfected WT added). (F) Rg4 CD8 T-cell recognition of the Mtb-infected macrophages in the antigen transfer assay. (G) Rg4 recognition in supernatant transfer assay. Unfiltered culture supernatants [SUP] from Mtb-infected macrophages were added to uninfected or macrophages infected using the indicated MOIs for 24-hours. IFN-γ production was measured as an indication of T-cell activation. Bars, mean ± SD. Data are representative of two (B - D), three (F) or four (G) independent experiments. Statistical testing: Spearman and linear regression test (D); two-way ANOVA with Dunnett’s test (F); or two-way ANOVA with Šidák correction for multiple comparisons (G). p values: *, p<0.05; **, p<0.01; ***, p<0.001; #, p<0.0001.

We next wished to determine the relative contributions of direct presentation by infected macrophages versus indirect presentation by bystander macrophages that secondarily acquired Mtb antigens through phagocytosis. An antigen transfer assay was established to mimic what might occur in the lung, where infected macrophages are surrounded by uninfected myeloid cells. MHC I^−/−^ macrophages were used as a control since they are unable to present antigens to CD8 T-cells. WT or MHC I^−/−^ macrophages were infected for 24 hours, and extensively washed to remove any extracellular bacilli. Then, uninfected WT or MHC I^−/−^ macrophages and TB10.4-specific CD8 T-cells were added, and IFN-γ was measured after 24 hours [Fig.5E]. To establish the baseline of non-cognate activation of the CD8 T-cells, MHC I^−/−^ macrophages and T-cells were added to infected MHC I^−/−^ macrophages [“K/K,” Fig.5F]. As the MOI was increased, a small but significant amount of IFN-γ was produced compared to the uninfected condition (i.e., MOI = 0). To determine the degree of direct presentation by primary-infected macrophages, uninfected MHC I^−/−^ macrophages and T-cells were added to infected WT macrophages. Under these conditions, IFN-γ increased as the MOI was increased, but did not become statistically significant compared to the “K/K” group until an MOI=8 [“W/K,” Fig.5F]. In contrast, the greatest amount of IFN-γ was produced when uninfected WT macrophages were added to infected cells, regardless of their genotype of the primary infected macrophage (i.e., WT or MHC I^−/−^) [see “W/W” and “K/W,” Fig.5F]. These results suggest that during high MOI infection, antigen transfer to bystander cells leads to indirect presentation to CD8 T-cells.

Other investigators have shown that soluble antigen, excreted antigen, or antigen-laden vesicles released by infected macrophages contain Mtb antigen and can be taken up by bystander macrophages and presented to CD8 T-cells^52–54^. To test this possibility, supernatants from macrophages infected overnight were collected. Addition of these unfiltered supernatants neither conferred nor enhanced presentation by uninfected or infected macrophages, respectively [Fig.5G]. Similar results were obtained with Mtb-infected macrophage cultures that were freeze-thawed to mimic cell necrosis. Although free antigen or vesicles contained in the supernatants could have diminished bioactivity secondary to dilution, our results suggest that efficient antigen transfer requires intact cells. We envision that engulfment of dying infected cells concentrates antigen or targets its delivery to a presentation-competent compartment^45,55^. Based on these data, we conclude that indirect presentation by uninfected macrophages could enhance CD8 T-cell activation during Mtb infection.

### The ESX-1 type VII secretion system is required for presentation of TB10.4 to CD8 T cells

Our finding that CD8 T-cell activation requires high MOI infection and is associated with macrophage death or occurs when uninfected macrophages secondarily acquired Mtb antigens, suggests that cell death of infected macrophages is a prerequisite for presentation of Mtb antigens to CD8 T cells. As the ESX-1 type VII secretion system is a major virulence factor that disrupts the phagosomal membrane and induces cell necrosis^56,57^, we hypothesized that ESX-1 would be required for MHC I presentation by macrophages. Therefore, we compared CD8 T-cell recognition of macrophages infected with wild-type H37Rv and a Mtb mutant with a deleted RD1 locus (ΔRD1), which removes most of the genes required for ESX-1 function^57,58^. Importantly, the EsxH gene that encodes the TB10.4 antigen is not located within the ESX-1 locus. Macrophages infected with the ΔRD1 mutant or H37Rv had similar intracellular bacterial burdens [Fig.6A, Left]. However, macrophages infected with the ΔRD1 mutant had little or no cell death compared to H37Rv-infected macrophages, either at 24 hours (i.e., when T-cells are added for the assay) or at 48 hours (i.e., when IFN-γ is measured) [Fig.6A, center and right]. The inflated viability of ΔRD1-infected macrophages results from staining of Mtb by crystal violet.

**Figure 6.**
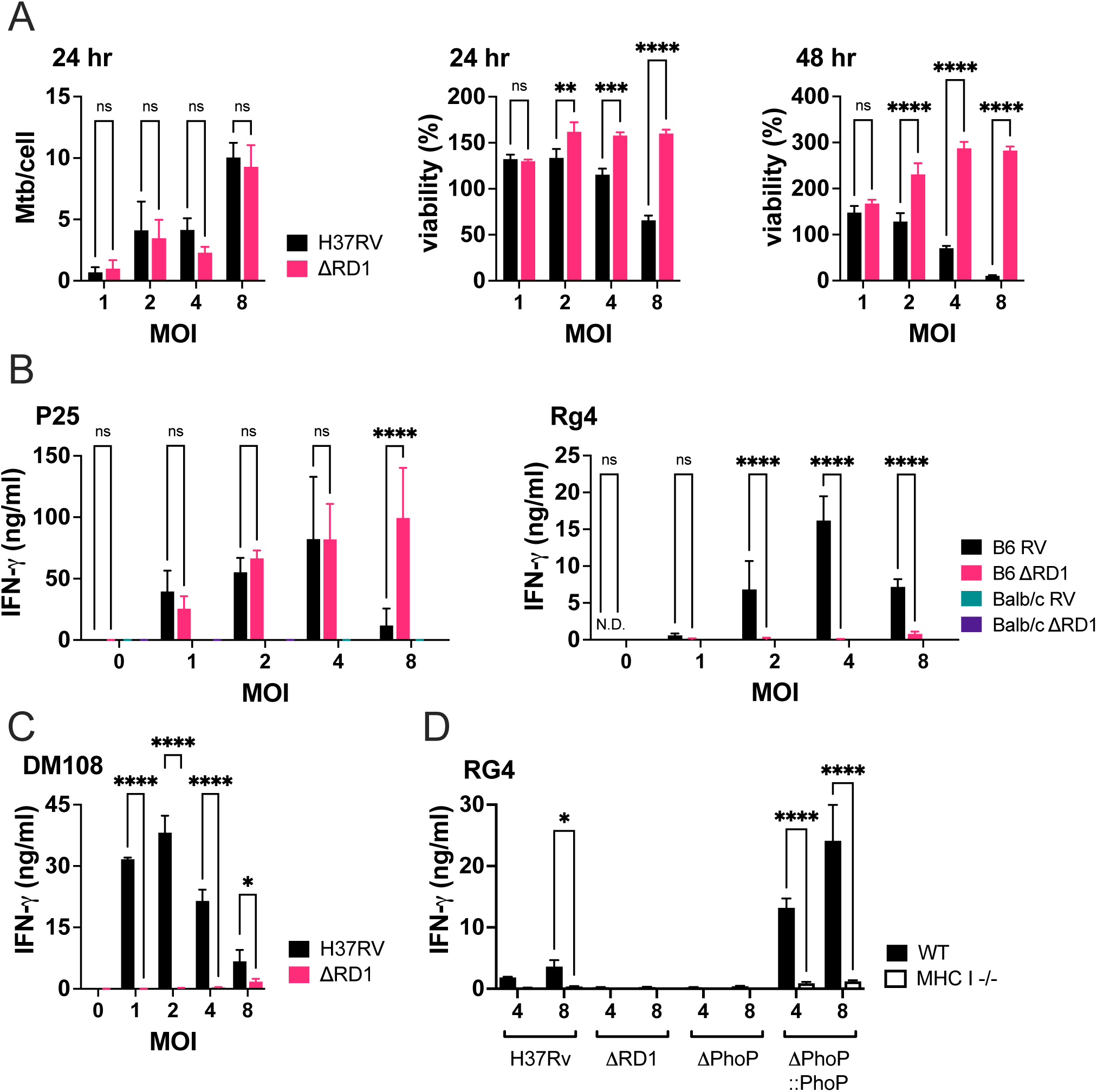
Cross-presentation is ESX-1 dependent. (A) Adjusted MOI (left) and normalized viability of HOXB8 macrophages infected overnight at the indicated MOIs with WT H37Rv or ΔRD1 Mtb at 24 hours (center) or 48 hours (right) post-infection (B) P25 (left) and Rg4 (right) recognition of H37Rv or ΔRD1 Mtb-infected B6 (matched) or BALB/c (mismatched) TGPM. (C) DM108 recognition of H37Rv or ΔRD1 Mtb-infected HOXB8 macrophages. (D) Rg4 recognition of WT or MHC I^−/−^ macrophages infected with H37Rv, ΔRD1, ΔPhoP or ΔPhoP-complemented strains of Mtb. T-cell activation measured by IFN-γ ELISA. Bars, mean ± SD (n=3 replicates/condition). Data are representative of four (A), three (B, C) or two (D) independent experiments. Statistical testing: two-way ANOVA with Šidák correction. p values: *, p<0.05; **, p<0.01; ***, p<0.001; #, p<0.0001.

Both ΔRD1- and H37RV-infected macrophages were recognized by Ag85b-specific CD4 T-cells [Fig.6B, left]. Interestingly, IFNγ-production by P25 CD4 T-cells was diminished at the highest MOI of H37Rv-infected but not ΔRD1-infected macrophages, which we suggest is due to APC death induced by fully virulent Mtb. In contrast, infection with ΔRD1, even at high MOIs, failed to activate two different TB10.4-specific CD8 T-cell lines [Fig.6B, 6C]. As the ΔRD1 mutant is difficult to complement and may have other defects, we validated our results by disrupting the PhoPR regulon (ΔPhoPR), which is required for the expression of the ESX-1 secretion system in H37Rv^48,59–61^. TB10.4-specific CD8 T-cells did not recognize ΔPhoPR-infected macrophages [Fig.6D]. In contrast, TB10-specific CD8 T-cells were activated by macrophages infected with the ΔPhoPR::PhoPR complemented strain. Thus, ΔPhoPR phenocopies the ΔRD1 strain and can be complemented. These data show that a high MOI is not sufficient for MHC I presentation; ESX-1 type VII secretion system is also required for presentation of TB10.4 by MHC I.

## Discussion

These studies expand upon our observation that TB10.4-specific CD8 T-cells poorly recognize Mtb-infected macrophages and extend the observation of inefficient MHC I antigen presentation to additional epitopes recognized by CD8 T-cells. Importantly, all the diverse types of macrophages used in this study can cross-present non-viable antigen to CD8 T-cells. Our results confirm that Mtb does not globally inhibit MHC I antigen presentation ^26^; instead, our data points to a defect specific to mycobacteria-containing compartments. Importantly, we do not believe that the poor recognition of infected macrophages is a feature only of the CD8 T-cell lines used in this study: only 10% of polyclonal CD8 T-cells from the lungs of infected mice produce IFN-γ when cultured with Mtb-infected macrophages ^22,41^.

We identify conditions that permit more efficient MHC I presentation of Mtb antigens to CD8 T-cells. *In vitro* infection of macrophages for 24 hours led to significantly more intracellular Mtb than a standard two-hour infection. More highly infected macrophages gain the capacity to present MHC I-restricted Mtb epitopes to CD8 T-cells. These data are consistent with work from the Lewinsohn lab showing that a CFP10-specific human CD8 T-cell clone preferentially recognizes highly infected human DCs ^62^. We confirm their original observation, extend it to macrophages, provide quantitative data, and show that this feature of host-pathogen interaction is conserved between species. We rule out the possibility that measuring only IFNγ underestimate T cell recognition of infected macrophages and address the role of noncognate activation and macrophage cell death. In contrast to CD8 T-cells, CD4 T-cells are activated even at a low MOI. Why is a high intracellular Mtb burden required for MHC I antigen presentation? One might assume that more bacilli lead to more antigen. However, we do not believe that antigen quantity is the key factor. We previously showed that overexpression of TB10.4 does not enable CD8 T-cell recognition of macrophages infected at low MOIs ^23^. Also, if this were only an issue of antigen abundance, high MOI infections of macrophages with Mtb strains containing deletions of the RD1 locus or the PhoP gene should be recognized by TB10.4-specific CD8 T-cells.

A high bacterial burden promotes cell death primarily through necrosis ^44,46–49^. We and others have shown that macrophages engulf dying infected cells ^45,63,64^. As DC can engulf dead, antigen-bearing cells and cross-present their antigens ^65–68^, we explored the possibility that the uptake of dying infected cells acted as a source of Mtb antigens that bystander macrophages could present via MHC I. Indeed, bystander macrophages acquired antigen from highly infected cells and presented TB10.4 antigen to CD8 T-cells more efficiently than primary infected macrophages. Although our data suggests that indirect presentation is the dominant pathway that activates CD8 T-cells, we cannot exclude the possibility that primary-infected macrophages also engulf dying infected macrophages and cross-present the bacterial antigens. Cell death directly contributes to high bacterial burden by facilitating bacterial growth, and reuptake by bystander macrophages, creating a cycle of increasing Mtb burden and necrosis ^50,51^. We cannot rule out that direct antigen presentation occurs. In particular, the cell interaction assay suggests that there is a correlation between T cell interaction with macrophages as a function of bacterial burden. In this assay, we cannot distinguish between macrophages that were infected with a great number of Mtb vs. those that acquired the bacilli secondarily. Furthermore, we hypothesize that indirect presentation requires direct contact with dead/dying cells since we did not observe cross-presentation after supernatant transfer. Thus, efferocytosis or phagocytosis may engulf dying infected cells, leading to the concentration of Mtb antigens, which are shuttled to cross-presentation-competent endosomes by scavenging receptors as has been shown for DNGR1 in DC ^67–69^.

The ESX-1 type VII secretion system, which is encoded within the RD1 locus, is one of the key virulence factors of Mtb and can cause phagosomal membrane damage ^56,57,70^. We show that ESX-1 is necessary for MHC I presentation of TB10.4 antigen by infected macrophages. While we believe that ESX-1-associated macrophage death is the major pathway that licenses cross-presentation, there are alternative hypothesis to consider. ESX-1 mediated phagosomal membrane damage could enable Mtb antigens access to the macrophage cytosol and facilitate MHC I presentation to CD8 T-cells. A high bacillary burden could be required for sufficient production of ESX-1 complexes to make the phagosomal membrane permeable to Mtb antigens. Cytosolic Mtb is sensed by cytosolic surveillance pathways including STING ^34,71^, which may augment cross-presentation of Mtb antigens. Finally, we favor the idea that activation of cell death pathways of Mtb-infected macrophages, leads to their engulfment by bystander macrophages. Skewing of cell death towards apoptosis will lead to efferocytosis and is associated with enhanced CD8 T-cell responses ^44,46,72^. While some macrophages infected with virulent Mtb will undergo apoptosis, many more are thought to undergo necrotic cell death. In this study, we did not define the mode of cell death, as both apoptotic and necrotic cells are engulfed by phagocytes ^64,67,73,74^. Interestingly, we previously reported that ESX-1 is not required for in vivo priming of TB10.4-specific CD8 T-cells ^75^. Also, a human Mtb8.4-specific CD8 T cell clone recognized Mtb-infected autologous DC independently of RD1 ^76^. Clearly, the abundance of Mtb proteins and how they are secreted will influence how they are sampled by the MHC I pathway. The disparity between the requirements for initial CD8 T-cell priming and recognition of infected macrophages at sites of infection could reflect that in vivo, death of macrophages by apoptosis (more frequent in the absence of ESX-1) and necrosis (more frequent in the presence of ESX-1) both ultimately lead to acquisition of Mtb antigens by APC.

The observation that CD8 T-cells respond to highly infected macrophages sheds light on our previous *in vivo* observations. It is paradoxical that TB10.4-specific CD8 T-cells don’t recognize infected macrophages in vitro but are a dominant and persistent populations in the lungs of infected mice. There is a discrepancy in the capacity of TB10.4-specific CD8 T-cells to mediate protection. Vaccination with TB10.4_4-11_ peptide does not protect mice against Mtb challenge, nor do TB10.4-specific CD8 T-cells enhance the ability of infected macrophages to control Mtb replication ^20,23^. However, Mtb-infected TCRα^−/−^ mice control Mtb infection better after transfer of TB10.4-specific CD8 T-cells and protection is TAP1-dependent ^21^. We surmise that in immunodeficient mice, the bacterial burden becomes so high that the transferred CD8 T-cells are activated by indirect cross-presentation, which as the main source of IFN-γ, protects the lungs.

If indirect presentation activates Mtb-specific CD8 T-cells in the lung during active TB, MHC I presented antigens could act as decoy antigens. CD8 T-cell recognition of Mtb antigens presented by uninfected APC could promote inflammation and promote Mtb transmission. In this scenario, the activated CD8 T-cells would have little impact on the outcome of infection. On the other hand, mediators (e.g., IFN-γ and TNF) released by CD8 T-cells activated by indirect presentation could activate nearby infected macrophages. CD8 T-cells might be important in mediating protection when there is significant bacterial growth and other immune mechanisms are failing. Recent work in rhesus macaques shows that CD8 T-cells contribute to protection late during infection ^77^ and CD8 T-cells are associated with granulomas that control Mtb infection in cynomolgus macaques ^78^. Pre-existing (i.e., vaccine-elicited) Mtb-specific CD8 T-cells could also play a role early during infection when the intracellular bacillary burden can exceed 20 bacilli/monocytic cells in the lung ^49^. However, our data predicts that this function could be equally performed by CD4 T-cells. An important issue to consider is whether certain antigens are more likely to be directly presented by Mtb-infected macrophages, as such antigens might be more effective in focusing the CD8 T-cells response on infected macrophages instead of uninfected bystander cells that have acquired Mtb antigens. We predict that developing approaches to distinguish Mtb antigens that are directly presented by infected macrophages from those that are indirectly presented would empower novel target selection for vaccines that elicit protective CD8 T-cells.

## Methods and Materials

### Ethics Statement

Studies were conducted using the relevant guidelines and regulations and approved by the Institutional Animal Care and Use Committee at the University of Massachusetts Chan Medical School (UMMS) (Animal Welfare A3306-01), using the recommendations from the Guide for the Care and Use of Laboratory Animals of the National Institutes of Health and the Office of Laboratory Animal Welfare.

### Aerosol Mtb Infection of Mice

Mice were infected by the aerosol route as previously described (Lee et al., 2020). Frozen bacterial stocks were thawed and diluted into 5 mL of 0.01% Tween-80 in PBS, then sonicated to create a bacterial suspension. Mice were infected with ~100 CFU Mtb Erdman via the aerosol route using a Glas-Col airborne infection system.

### Intravascular Staining

Mice were injected intravenously with 2.4 µg/mouse of anti-CD45 – BV750 in 200 µL of injection medium (2% FBS in PBS) 2 minutes before euthanizing with CO_2_. Lungs were then perfused and removed 3 minutes after anti-CD45 injection. Lymphocytes from blood collected in RPMI containing 40 U/mL of heparin were isolated using Lympholyte (CEDARLANE) and analyzed by flow cytometry to confirm uniform staining with anti-CD45 – BV750.

### Generation of single-cell suspensions from tissues

Lung single-cell suspensions from Mtb infected mice were isolated as previously described (Lu et al., 2021). Briefly, mice were euthanized with CO_2_ and lungs were perfused with 10 mL RPMI (GIBCO) before harvest. Single cell suspensions were prepared by homogenizing lungs using a GentleMACS tissue dissociator, digesting with 300 U/mL collagenase (Sigma) in complete RPMI (10% FBS, 2mM L-Glutamine, 100 U/mL Penicillin/Streptomycin, 1 mM Na-Pyruvate, 1X non-essential amino acids, 0.5X Minimal essential amino acids, 25 mM HEPES, and 7.5 mM NaOH) at 37C for 30 minutes, and followed by a second run of dissociation using the GentleMACS. Suspensions were then filtered through 70 µm strainers and incubated with 1 mL ACK Lysis Buffer (GIBCO) for 1 minute at room temperature (25C). The cells were resuspended in suitable media or buffer for further use and filtered through 40 µm filters.

### Macrophages and DC

Thioglycolate elicited peritoneal macrophages (TGPM) were harvested by first injecting mice with 2 ml of 3% thioglycolate broth intraperitoneally. 4-5 days after injection, macrophages were extracted through peritoneal lavage and purified with Miltenyi Biotec CD11b microbeads (Catalog #130-049-601). HOXB8 progenitor cells were established as described ^79^. MHC I KO lines were generated with bone marrow from the β2 microglobulin knockout line B6.129P2-B2mtm1Unc/DcrJ (Jackson). Progenitor cells were collected, washed twice with PBS, and cultured in 150 x 15 mm non-adherent plates in the appropriate media. For macrophages, progenitors were cultured in 20% L929 media for 8-12 days before adherent cells were harvested. DC were cultured in either 10% GM-CSF or 5% FLT3 as indicated in the results. BMDC were prepared as previously described ^80^. Briefly, bone marrow harvested from murine femurs was washed and cultured in 10% GM-CSF complete media for 10 days. Non-adherent cells were harvested and used for infections.

### Protein-coated beads

BioMag®Plus Amine protein coupling kit (Bangs Laboratories Inc.) (Catalog #86000-1) was used to conjugate full length Ovalbumin (Sigma-Aldrich #A5503) or TB10.4 (manufactured by Genscript) protein to magnetic beads according to the manufacturer’s protocol. Briefly, BioMag®Plus Amine particles were washed, activated, and magnetically separated before being incubated with protein for conjugation. The beads were further washed, and the reaction was quenched before being diluted for use in assays.

### Infections

*Mycobacterium tuberculosis* strains were cultured in Middlebrook 7H9 broth until an OD_600_ of 0.8-1.2 was reached. Cultures were washed by centrifugation and incubated in TB coat (1% FBS, 2% human serum and 0.05% Tween-80 in RPMI) for 5 minutes to opsonize bacteria and impede clumping. Bacteria were washed again and resuspended in complete media without antibiotics. Suspensions were filtered through a 5 μm filter to remove clumps and absorbance was measured to estimate bacterial density. Diluted Mtb was added to target cells at the corresponding MOI and left for 2 (low MOI infections) or 18 – 24 (high MOI infections) hours at 37°C. Cells were then washed 3-4 times with warm RPMI to remove extracellular bacteria.

### ΔphoPR mutant and complementation

The ΔphoPR mutant of M. tuberculosis H37Rv was isolated following allelic exchange of 2.178 kb region of the phoP-phoR (rv0757-rv0758) sequence with a hygromycin-chloramphenicol cassette as described previously (Lee et al., 2011). The hygromycin-resistant ΔphoPR mutant retained the first 32 bases of phoP gene and the last 36 bases of the phoR gene. For complementation, a 3.3 kb DNA fragment containing the wildtype allele of H37Rv phoPR coding sequence and 1 kb upstream region encompassing the putative promoter was amplified by PCR and cloned into the kanamycin mycobacterial L5 attB integrating vector pMV306 (MedImmune Inc., Gaithersburg, MD) to construct the phoPR complementing plasmid pKP406. Following sequence verification of the cloned insert, the KanR plasmid pKP406 was electroporated into the hygromycin-resistant M. tuberculosis ΔphoPR mutant and the transformants were selected on 7H10 plates supplemented with 50 μg/ml hygromycin and 25 μg/ml kanamycin.

### Polyclonal T-cell lines

Mice were vaccinated using the Trivax vaccination strategy^81^ with either TB10.4_20-28_ or Mtb32A_309-318_ two to three times, three weeks apart, as previously described^20^. The frequency of antigen specific CD8 T-cells was estimated by flow cytometry after staining peripheral blood with the Mtb-specific tetramers K^b^/Mtb32A_309-318_ and K^d^/ TB10.4_20-28_ provided by the NIH Tetramer Core. Mice with positive vaccine responses were euthanized to allow collection of the spleens and peripheral lymph nodes. CD8 T-cells were isolated using Stem Cell CD8 isolation kits (Catalog #19753). Purified CD8 T-cells were cultured with peptide-pulsed irradiated splenocytes to activate and expand antigen-specific CD8 T-cells. The polyclonal cells were successively restimulated using irradiated splenocytes, peptide, and IL-2, every 4 weeks to expand and enrich antigen-specific populations. Antigen-specific CD8 T-cells were quantified using tetramers supplied by the NIH tetramer core.

### T-cell recognition assays

Macrophages were seeded in 96 well float bottom TC-treated plates at a density of 100,000 cells/well and infected as described above. T-cells were added to infected macrophages in a 1:1 ratio unless noted otherwise. For ELISA analysis, supernatants were collected after co-culturing the cells for 24 hours at 37°C. Supernatants were filtered through 0.2 μm filter plates to exclude any bacteria, and cytokine levels were quantified with Biolegend ELISA kits (Catalog #430801). For flow cytometry, T-cells were co-cultured with infected macrophages for 6 hours at 37°C with brefeldin A (Golgiplug). Cells were then stained for surface markers and intracellular cytokines as described ^10^.

### Immunofluorescence microscopy staining

T-cells were cocultured with infected macrophages for the time indicated. Samples were washed with PBS and fixed with 4% paraformaldehyde for 20 minutes. Samples were washed with PBS and blocked in 1% BSA for 30 minutes at room temperature. Samples were then washed and incubated in the appropriate primary antibodies overnight at 4°C. Cells were then washed and incubated in secondary antibody for one hour at room temperature.

### Fluorescent beads for confocal microscopy

We used Polysciences Fluoresbrite® YG Microspheres 3.00µm for confocal imaging analysis. Beads were washed in PBS with 1% BSA, then opsonized with TB coat (1% FBS, 2% human serum and 0.05% Tween-80 in RPMI) at room temperature for 30 minutes to facilitate phagocytosis. Beads were diluted to 5 x 10^6^ beads/ml and added alongside H37Rv Mtb for cell infections.

### Confocal imaging and analysis

Images were acquired on a Leica SP8 confocal microscope at on an Apo TIRF 60x Oil DIC N 2 objective with a numerical aperture of 1.49. We used 405, 488 and 640 nm lasers, and 450/50 nm and 525/50 emission filters. Pixel size is 207 x 207 nm, and the voxel depth is 400 nm. 18-24 Z-steps were acquired for each image with a step size of 0.4 μm. Colocalization of targets with Mtb were quantified with ImageJ by calculating Mander’s coefficients (Fraction of overlapping pixels between two channels) of the signal of interest with Mtb YFP signal. Regions of interests (ROIs) were manually selected around individual bacilli before coefficients were calculated to quantify the amount of colocalization of our target with Mtb on a cell-cell basis.

### Slide scanning Immunofluorescence

Images were acquired with a TissueFAXS iQ tissue cytometer (TissueGnostics GmbH) built on a Zeiss AxioImager.Z2 microscope base with Märzhäuser stage, Lumencor Spectra III light engine, Zeiss EC Plan-Neofluar 20x/0.75 NA objective and Hamamatsu Orca Fusion BT C15440 camera. DAPI filters were 390/22 excitation, 89402 quad dichroic, 89402m emission. AF488 filters were 475/20 excitation, T495lpxr dichroic, ET519/26 emission. TRED filters were 555/28x excitation, 89402bs dichoric, and 89402m emission. Cy5/AF647 filters were 631/28x excitation, 89402bs dichoric, and 89402m emission. AF740/Cy7 filters were 747/11 excitation, T770lpxr and ET810/90m emission.

### Strataquest Image analysis and quantification

Whole slide scans were analyzed with Tissuegnostics Strataquest Plus V7.1.1.127 software (TissueGnostics GmbH). We developed analysis pipelines to identify and differentiate T-cells and macrophages based on nuclei identification, staining intensity of F4/80 and CD3, and a cellular mask calculation. Once T-cells and macrophages were identified, we extrapolated total YFP signals for each macrophage to calculate approximate bacterial burdens. We converted the total YFP signal to Mtb numbers by dividing the signal by the average signal of one Mtb bacilli. Next, we also calculated the number of T-cells in contact with each macrophage to correlate Mtb burden with T-cell contacts. Statistical testing including correlations between Mtb burden and T-cell contacts were calculated in GraphPad Prism. We binned macrophages based on their Mtb burden with a bin size of one, then calculated and plotted the mean T-cell contacts of each bin. Between 3,100-6,900 macrophages were analyzed per condition. MOIs greater than 24 were not analyzed because there were generally fewer than 5 cells in these bins. In contrast, an average of 34 macrophages were analyzed at each MOI between 4-14.

### ATP Luciferase viability assay

Cells were infected with Mtb as described above. We used the Abcam Luminescent ATP Detection Assay Kit (ab113849) and followed the provided protocol. Briefly, cells were lysed with detergent for five minutes to release ATP. Substrate was added and incubated in the dark for 15 minutes. Luminescence of the wells was measured, and samples were compared to standards to determine cell viability.

### Crystal Violet cell viability

Mtb-infected cells were washed and fixed with 4% paraformaldehyde for 20 minutes at room temperature. Cells were then washed with PBS and incubated with 0.1% crystal violet in water for 30 minutes. Plates were washed by submerging the plate in a container of fresh tap water 3-4 times. Cells were then lysed with 150 μL of 0.2 % Triton X-100 in water for 30 minutes to release crystal violet. Samples were mixed and transferred to a new flat-bottom clear plate and absorbance was measured at 600 nM. Viability was determined as follows: %Viability = ((Ab600 Control – Ab600 Experimental) / Ab600 Control))*100.

### CFU determination

Infected macrophages were washed 3X with warm cRPMI to remove extracellular bacteria. Cells were then lysed with 1% Triton X-100 in PBS for 2 minutes. Samples were serially diluted and plated on 7H11 Agar plates from Hardy Diagnostics (Catalog #W40). Plates were incubated at 37°C for 21 days when colonies were counted and Mtb/well was calculated based on the dilutions.

### Adjusted MOI calculations

To account for cell death in our Mtb/cell calculations, we calculated the average number of viable cells per condition by multiplying %viability by the number of cells seeded. If the %viability was quantified at over 100%, we adjusted it to 100%. The number of bacteria/well determined by CFU calculations was divided by the number of viable cells to determine approximate Mtb/cell numbers (# of Mtb / # of cells = Mtb/cell).

### Antigen Transfer Assay

B6 WT or MHC I KO macrophages were seeded in 96 well flat bottom tissue culture-treated plates at a density of 50,000 cells/well and infected as described above. Following the infection, 50,000 fresh WT or MHC I KO macrophages were added to the wells along with 100,000 T-cells. Co-cultures were left at 37°C for 16-24 before samples were collected for analysis. Prior to infection, primary-infected macrophages were stained with CellTracker™ Violet BMQC (VioGreen channel) (ThermoFisher # C10094) according to the manufacturer’s protocol. The macrophages were infected as described above using the 24-hour (“high MOI”) protocol. After extensive washing to remove any remaining extracellular bacteria, bystander (uninfected) macrophages, which had been stained with CellTracker™ Deep Red (ThermoFisher # C34565) to differentiate them from the primary infected macrophages, were added to the culture. The macrophages were cultured for 24 hours, then they were washed and analyzed by flow cytometry or fluorescent microscopy. For microscopy, samples were imaged with the slide scanning software and analyzed with strataquest as described above. Correlations between primary dye and Mtb were calculated and plotted using GraphPad Prism.

## Acknowledgments

We thank the Sanderson Center for Optical Experimentation (SCOPE) core facility at UMass Chan Medical School (Worcester, MA), which provided access to the Nikon A1 confocal and the TissueGnostics TissueFAXS SL tissue cytometer. The SCOPE TissueFAXS SL was funded by a Massachusetts Life Science Center Bits to Bytes award to Drs. Christina Baer and Dorothy Schafer. The TB10.4 and 32A tetramers were obtained from the NIH Tetramer Core Facility. This work was funded by NIH/NIAID grants R01AI106725 and R01 AI123286 to S.M.B. The funders had no role in the design or conduct of the study, nor the interpretation of the data and the preparation of the manuscript.

## Supplemental Figure Legends

**Figure S1. Markers of cross-presentation co-localize with Mtb containing compartments in infected macrophages**. TGPM were infected with H37Rv.YFP for two hours at an MOI of 5, followed by extensive washing to remove any extracellular bacilli. Two or twenty-four hours after infection, the macrophages were fixed and stained with antibodies specific for proteins known to have a role in cross-presentation by APC to CD8 T-cells. Colocalization between H37Rv.YFP and MHC I, P97, Rab5, or Cathepsin S was determined by confocal fluorescent microscopy and Mander’s coefficients were calculated for each infected macrophage. Each data point represents an individual macrophage. Bar, mean. Statistical analysis by one-way ANOVA with Fisher’s LSD test. p values: *, p<0.05; **, p<0.01; ***, p<0.001; #, p<0.0001

**Figure S2. Mtb-infected BMDC and HOXB8-differentiated DCs fail to cross-present TB10.4 to CD8 T-cells in vitro under conventional culture conditions.** (A) Rg4 CD8 T-cell recognition of H37Rv-infected BMDC. B10 or B10.BR BMDC (H-2-matched and mismatched [H-2^k^], respectively) were pre-activated with LPS (100 ng/ml) or not and pulsed with TB10.4_4-11_ peptide or infected with H37Rv at MOIs of 5 or 20 for two hours. After 24 hours, IFN-γ was measured as an indication of CD8 T-cell activation. (B) Rg4 recognition of H37Rv-infected HOXB8-derived DC. GM-CSF HOXB8 progenitor cells were cultured in either GM-CSF or Flt3 to generate DC, or in M-CSF to generate macrophages ^38,79,82^. The differentiated APC were pulsed with TB10.4_4-11_ peptide as a positive control or infected with H37Rv at an MOI of 5 for 2 hours. After 24 hours, IFN-γ was measured as an indication of CD8 T-cell activation. Bars, mean ± SD. Data are representative of two (A) or four (B) independent experiments, each with 3 technical replicates/condition. Statistical analysis by one-way ANOVA with Fisher’s LSD test. p values: *, p<0.05; ***, p<0.001; #, p<0.0001. N.D., not detected. Dotted line, upper limit of detection. Not all statistical comparisons are shown for clarity. UI, uninfected; P, peptide; INF, Mtb-infected.

**Figure S3. BATF3 is not required for priming of the immunodominant CD8 T cell responses to 32A or TB10.4.** Batf3 is required for the development of CD8α^+^ populations and optimal cross-priming of CD8 T-cells ^39^. We infected Batf3^−/−^ and wild-type B6 mice to compare Mtb-specific CD8 T-cell responses using specific tetramers. We found small but statistically insignificant decrease in 32A and TB10.4 CD8 T-cell responses 2 (A) or 4-6 (B) weeks post infection. n=5 mice/group. 4- and 6-week data were pooled form two experiments. Statistical testing by multiple t-tests. p values: *, p<0.05; **, p<0.01; ***, p<0.001; #, p<0.0001. Not all comparisons are shown for clarity.

**Figure S4. Excessive macrophage cell death is associated with reduced antigen presentation to CD4 and CD8 T-cells.** (A) B6 TGPMs were infected overnight with H37Rv at the indicated MOIs. After washing, Rg4 CD8 or P25 CD4 T-cells were added, and T-cell recognition of the infected macrophages was based on measurement of IFN-γ after 24-hours. In this experiment, P25 CD4 T-cells more sensitively recognized Mtb-infected TGPM at lower MOIs (i.e., MOI=1). However, neither the CD4 not the CD8 T-cells recognized Mtb-infected TGPM at the highest MOI (i.e., MOI=10). (B) Viability assays using Crystal Violet (left) or ATP-luciferase (right) of TGPMs 24 hours post H37Rv infection was performed using a set of macrophages infected in parallel with the macrophages used in Fig.S5A. Bars, mean±SD. Data representative of three (A) or two (b) independent experiments with 3-6 replicates/condition. Statistical analysis by one-way ANOVA, using the Dunnett test compared to uninfected macrophages. p values: *, p<0.05; **, p<0.01; ***, p<0.001; #, p<0.0001.

