## Supplementary figures and images for "High bacillary burden and the ESX-1 type VII secretion system promote MHC class I presentation by *Mycobacterium tuberculosis* infected macrophages to CD8 T-cells"

### Supplemental Figures 1-4

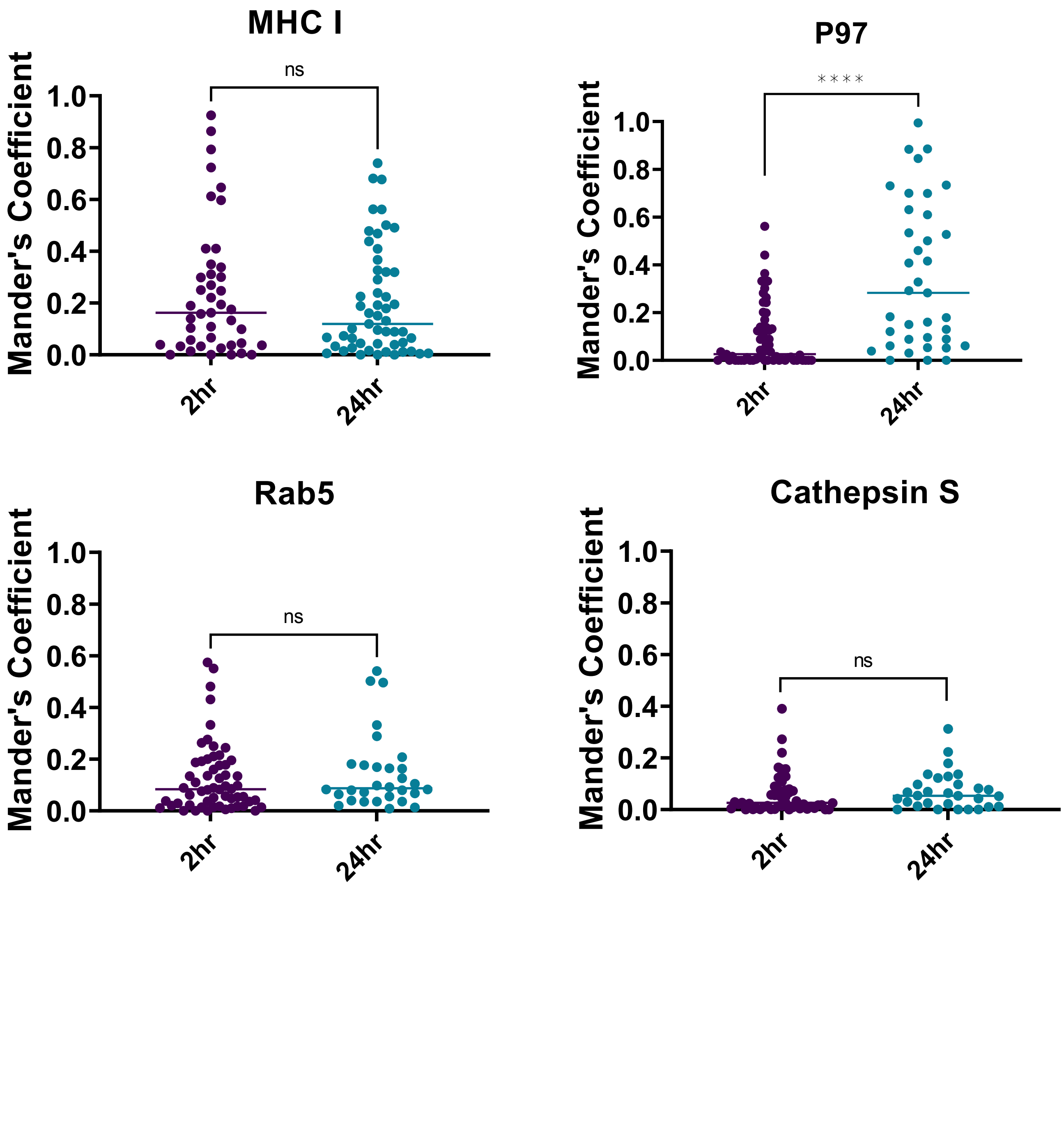

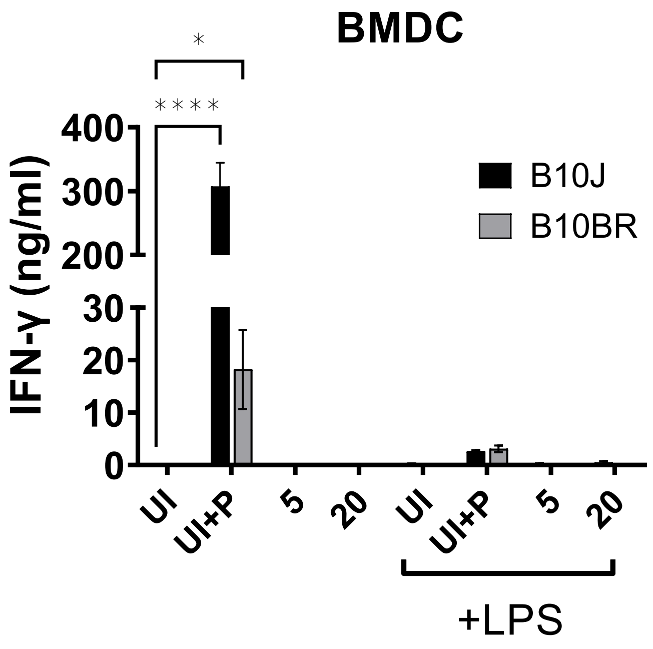

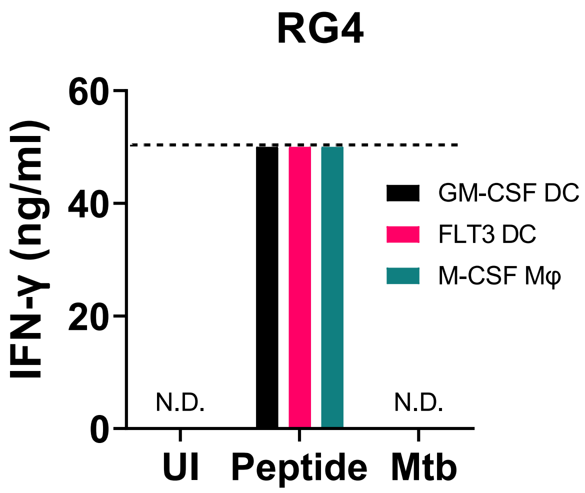

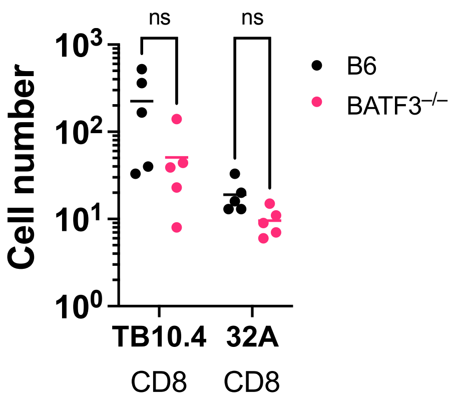

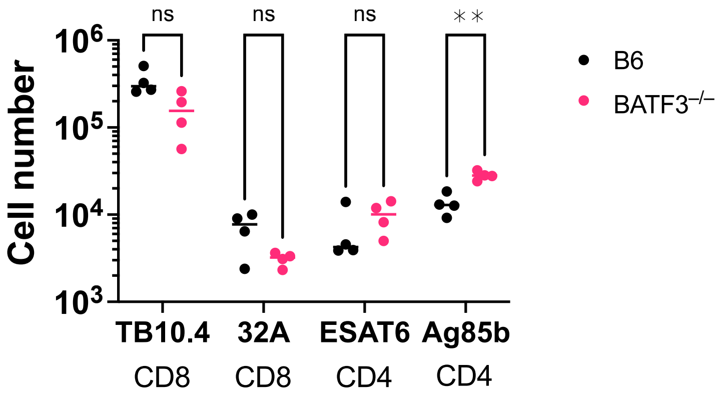

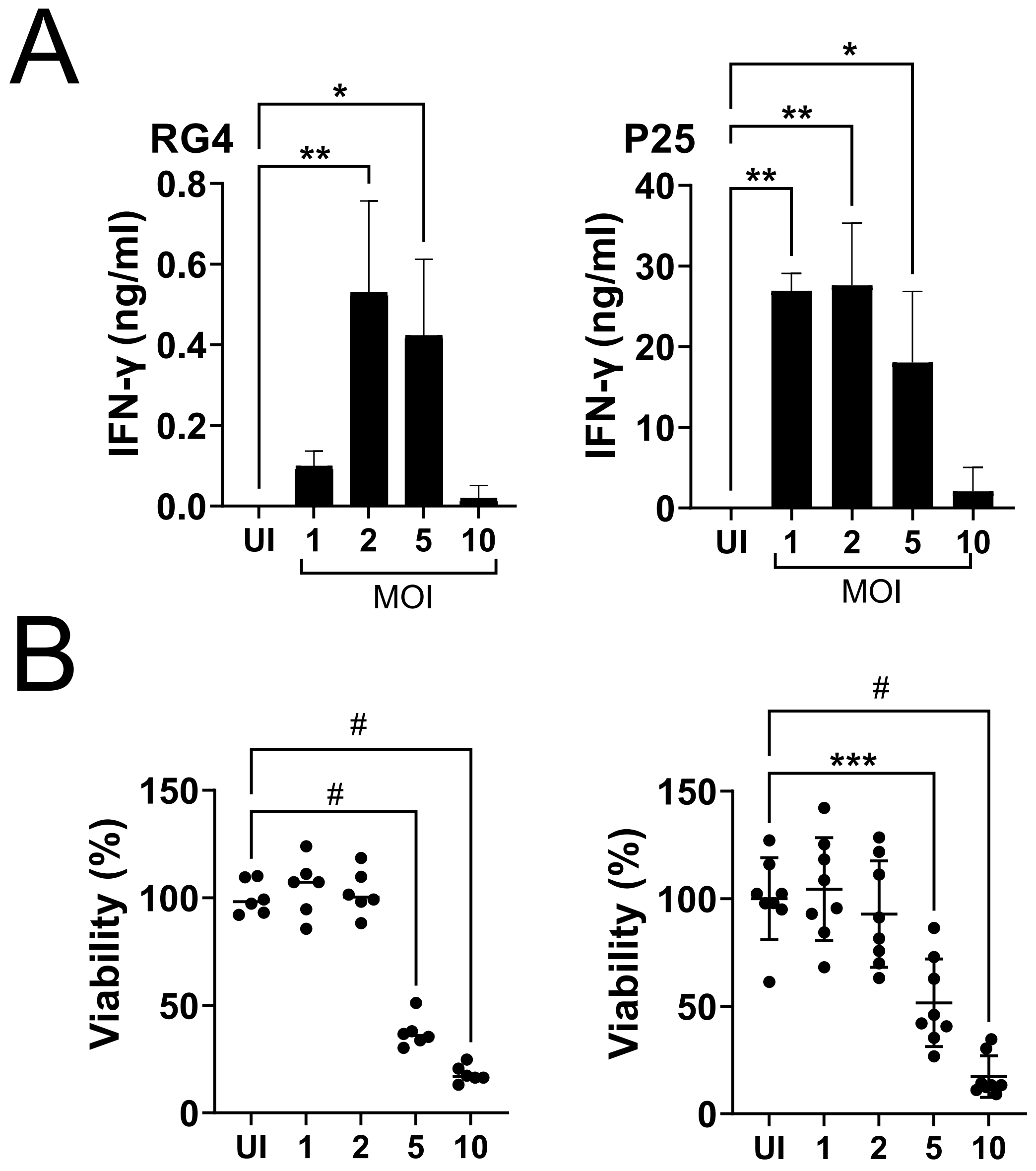
